# Ccr2 suppression by minocycline in Cx3cr1/Ccr2-visualized inherited retinal degeneration

**DOI:** 10.1101/2020.09.01.277285

**Authors:** Ryo Terauchi, Hideo Kohno, Sumiko Watanabe, Saburo Saito, Akira Watanabe, Tadashi Nakano

## Abstract

Retinal inflammation accelerates photoreceptor cell death (PCD) caused by retinal degeneration. Minocycline, a semisynthetic broad-spectrum tetracycline antibiotic, has previously been reported to show PCD rescue effect in retinal degeneration. The purpose of this study was to assess the effect of minocycline on Cx3cr1 and Ccr2 expression in retinal degeneration. *Mertk^-/-^Cx3cr1^GFP/+^Ccr2^RFP/+^* mice, which enabled observation of *Cx3cr1*- and *Ccr2*-expression pattern in inherited retinal degeneration, were used to test the effect of minocycline. Minocycline was systemically administered to *Mertk^-/-^Cx3cr1^GFP/+^Ccr2^RFP/+^* mice. For observing the effect of minocycline on Cx3cr1 and Ccr2 expression, administration was started on 4-week-old mice and continued for 2 weeks. To assess the PCD rescue effect, minocycline was administered to 6-week-old mice for 2 weeks. The expression pattern of Cx3cr1-GFP and Ccr2-RFP were observed on retinal and retinal pigment epithelium (RPE) flat-mounts. The severity of retinal degeneration was assessed on retinal sections. Minocycline administration suppressed *Ccr2* expression in *Mertk^-/-^Cx3cr1^GFP/+^Ccr2^RFP/+^* mice as observed in retinal and RPE flat-mounts. On the contrary, Cx3cr1 expression was not affected by minocycline administration. Retinal degeneration is ameliorated in minocycline administered *Mertk^-/-^Cx3cr1^GFP/+^Ccr2^RFP/+^* mice. In conclusions, Minocycline suppression of Ccr2 expression correlates to amelioration of retinal degeneration.

## Introduction

Inflammation in the central nervous system (CNS), as well as the retina, is considered as a complicating factor in degenerative diseases (1–5). Retinal inflammation is regarded to accelerate photoreceptor cell death (PCD) in retinal degeneration (RD), including age-related macular degeneration (AMD) and retinitis pigmentosa (RP) (6). Hence, management of inflammation is pivotal and presumably beneficial for patients with RD. Thus, the elucidation of inflammatory mechanisms for management of RD is a major research focus.

Minocycline, a semisynthetic, broad-spectrum tetracycline antibiotic, shows anti-inflammatory properties (7). Several studies, including ours, show that minocycline can ameliorate PCD in RD (8–10). However, the mechanism of PCD rescue effect by minocycline remains largely unknown. Two potential mechanisms have been suggested that include action through its anti-apoptotic properties and anti-inflammatory effect (11). The innate immune system, which has a rapid non-specific response to an antigen, has been implicated in the development of retinal degeneration including human AMD and RP (12). In healthy retina, microglia are located in the outer and inner plexiform layers and survey retinal homeostasis like guardians of the retina (12). However, in the stage of retinal degeneration, microglia get activated and migrate to outer retina and subretinal space, the space between the outer segments of photoreceptors and retinal pigment epithelium (RPE). As minocycline inhibits both microglial activation and migration, it is regarded that microglial suppression by minocycline is a major mechanism of PCD rescue (11). However, minocycline is not a microglia-specific drug. Furthermore, the other protagonist of retinal inflammation, bone marrow-derived macrophages, invade the outer retina (10, 13, 14). Therefore, the demonstration of why and how minocycline rescues photoreceptor cells in the degenerative stage is still important.

Recently, we developed *c-mer proto-oncogene tyrosine kinase* (*Mertk*)^-/-^*Cx3cr1^GFP/+^Ccr2^RFP/+^* mice, which enable the observation of Cx3cr1 and Ccr2 expression pattern in inherited retinal degeneration without requiring any non-physiological procedures, such as doxycycline administration (widely used for tetracycline-controlled transcriptional activation), or light damage (15). Before retinal degeneration occurs, only *Cx3cr1* expression is observed corresponding to resting microglia (16). In progressive retinal degeneration, Ccr2 expression is markedly increased (15). Due to this observation, we considered our model to be suitable for the elucidation of inflammatory targets and candidate drugs, such as minocycline.

In this study, we report that minocycline administration to *Mertk^-/-^Cx3cr1^GFP/+^Ccr2^RFP/+^* mice shows not only PCD amelioration but also suppression of *Ccr2* expression. The expression of *Ccr2* in the outer retina and subretinal space is reduced with minocycline administration. Taken together, *Ccr2* suppression is one of the mechanisms in photoreceptor cell rescue achieved through minocycline administration.

## Materials and methods

### Animals

The *Mertk^-/-^Cx3cr1^GFP/+^Ccr2^RFP/+^* mice were generated as previously described.(15) Genotyping for *Mertk* was performed with the following primers: wild type, forward 5’-GCTTTAGCCTCCCCAGTAGC-3’, reverse 5’-GGTCACATGCAAAGCAAATG-3’; mutant, forward 5’-CGTGGAGAAGGTAGTCGTACATCT-3’ and reverse 5’-TTTGCCAAGTTCTAATTCCATC-3’. Genotyping for *Cx3Cr1* was performed with the following primers: wild type, forward 5’-TCCACGTTCGGTCTGGTGGG-3’ and reverse 5’-GGTTCCTAGTGGAGCTAGGG-3’; and *Cx3cr1* mutant, forward 5’-GATCACTCTCGGCATGGACG-3’ and reverse 5’-GGTTCCTAGTGGAGCTAGGG-3’. Genotyping for *Ccr2* was performed with the following primers: common, forward 5’-TAAACCTGGTCACCACATGC-3’; wild type, reverse 5’-GGAGTAGAGTGGAGGCAGGA-3’; and *Ccr2* mutant, reverse 5’-CTTGATGACGTCCTCGGAG-3’.

Equal numbers of male and female mice were used. All mice were housed in the animal facility at the Jikei University School of Medicine, where they were maintained either under complete darkness or on a 12 h light (~10 lux)/12 h dark cycle. All animal procedures and experiments were approved by the Jikei University School of Medicine Animal Care Committees and conformed to both the recommendations of the American Veterinary Medical Association Panel on Euthanasia and Association for Research in Vision and Ophthalmology Statement for the Use of Animals in Ophthalmic and Vision Research.

### Minocycline administration

Minocycline was purchased commercially (Sigma-Aldrich, St. Louis, MO) and dissolved in PBS for administration via intraperitoneal (IP) injection.

### Flat-mount retina and RPE preparation

All procedures for retina and RPE flat-mounts were carried out as described previously (10). Images of flat-mounts were captured by a confocal microscope (LSM, Carl Zeiss, Thornwood, NY). For the retina flat-mount, the entire retina was captured at 5 μm intervals and all photographs were projected in one slice. For the RPE flat-mounts, the entire visible RPE was captured at 3 μm intervals and projected in one slice.

### Histological analysis

All retinal sections were prepared using previously described procedures (10, 17). Cx3cr1-GFP or Ccr2-RFP positive cell number was counted using ImageJ (National Institutes of Health, Bethesda, MD). Immunohistocytology images were captured by a confocal microscope (LSM 880, Carl Zeiss, Thornwood, NY).

### Data analysis

Data represent the mean ± SD. At least three independent experiments were compared using the one-way analysis of variance test.

## Results

### Characterization of *Mertk^-/-^Cx3cr1^GFP/+^Ccr2^RFP/+^* mice retina

In *Mertk^-/-^Cx3cr1^GFP/+^Ccr2^RFP/+^* mice, the descriptions “4-” and “6-week-old” correspond to the retinal non-degenerative stage and RD ongoing stage, respectively (15). Representative Cx3cr1-GFP single-positive cells observed in retinal flat-mount of 4-week-old *Mertk^-/-^Cx3cr1^GFP/+^Ccr2^RFP/+^* mice, and Cx3cr1/Ccr2 dual-positive cells observed in RPE flat-mount of 6-week-old *Mertk^-/-^Cx3cr1^GFP/+^Ccr2^RFP/+^* mice are shown in Fig. 1A and B. Time series vertical sections of *Mertk^-/-^Cx3cr1^GFP/+^Ccr2^RFP/+^* mice are shown in Fig. 1C-E. At “4-week-old”, only Cx3cr1 expression was visible in inner retina (Fig. 1C). Neither Cx3cr1 nor Ccr2 expression were observed in outer retina and subretinal space. At 3-month-old, retinal degeneration, represented by outer nuclear layer (ONL) thinning, was observed (Fig. 1D). The number of ONL nuclei decreased from approximately 12 (4-week) to 1-4 (3-month). Abundant Cx3cr1 and Ccr2 expression were observed in ONL and subretinal space (Fig. 1 D1-D4). Some of cells were Cx3cr1/Ccr2 dual-positive. At 1.5-year-old, almost all nuclei in ONL had diminished indicating severe retinal degeneration (Fig. 1E). The frequency of Cx3cr1 and Ccr2 expression observed was less (data not shown) compared to degeneration ongoing stage (e.g., from 6-week to 3-month). The expression of Cx3cr1 and Ccr2 observed part of 1.5-year-old *Mertk^-/-^Cx3cr1^GFP/+^Ccr2^RFP/+^* mice is shown (Fig. 1 E1-E4).

**Fig 1.**
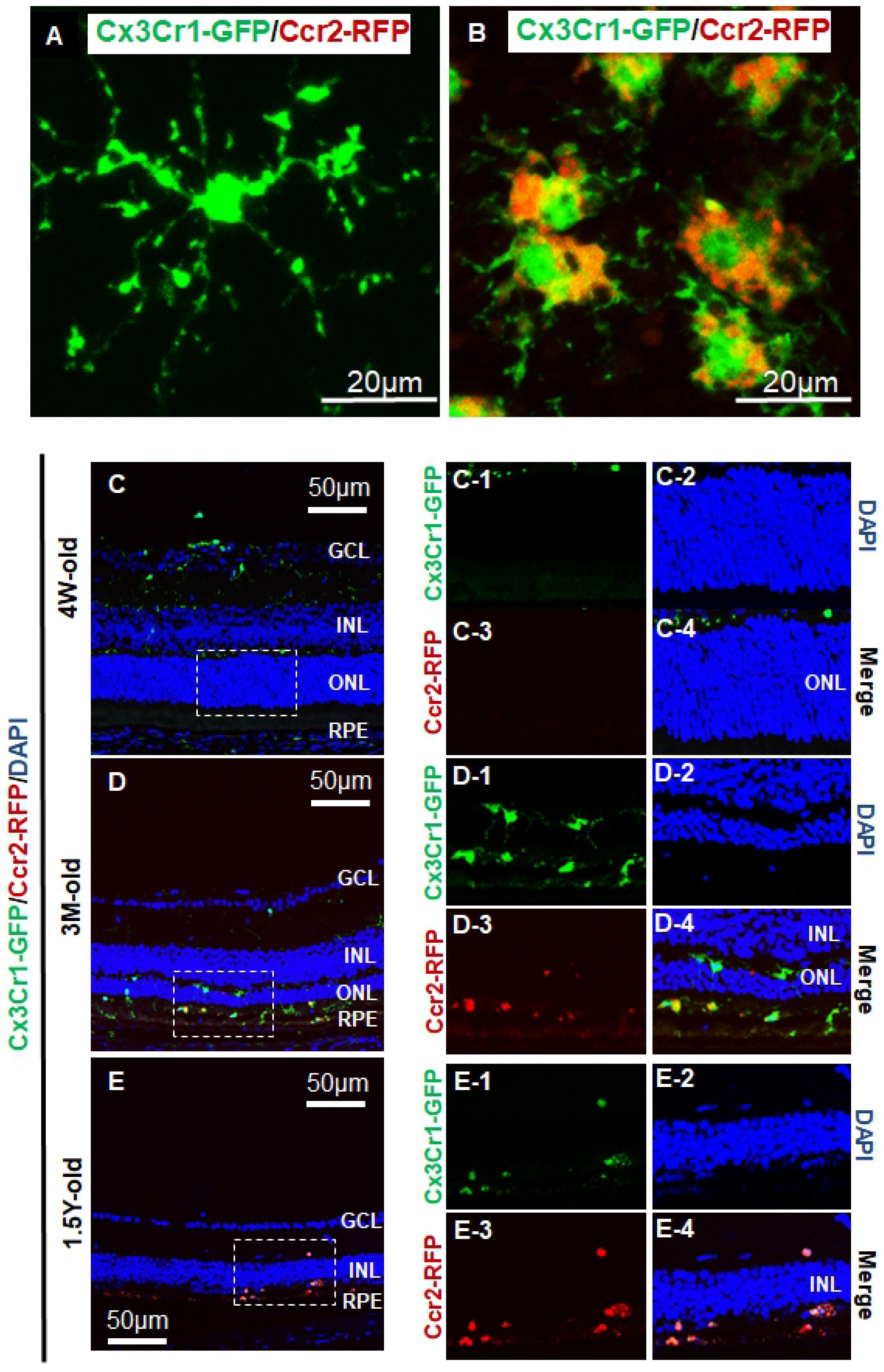
Characterization of *Mertk^-/-^Cx3cr1^GFP/+^Ccr2^RFP/+^* mice. Magnified Cx3cr1-positive cells observed in 4-week-old *Mertk^-/-^Cx3cr1^GFP/+^Ccr2^RFP/+^* mice retina flat-mount (A) and Cx3cr1/Ccr2 dual-positive cells in 6-week old *Mertk^-/-^Cx3cr1^GFP/+^Ccr2^RFP/+^* mice RPE flat-mount are shown. Vertical sections from 4-week, 3-month, and 1.5-year old *Mertk^-/-^Cx3cr1^GFP/+^Ccr2^RFP/+^* mice are shown (C-E). Inserts are shown as magnified images (C1-C4, D1-D4, E1-E4). GCL, ganglion cell layer; INL, inner nuclear layer; ONL, outer nuclear layer; RPE, retinal pigmented epithelium.

### Minocycline administration suppressed Ccr2 expression in neural retina

Minocycline was administered to *Mertk^-/-^Cx3cr1^GFP/+^Ccr2^RFP/+^* mice, between 4 and 6 weeks of age (continuous 14-day daily IP administration). The minocycline group was divided into 50 mg/kg (Mino50) or 100 mg/kg (Mino100) administration. Phosphate-buffered saline (PBS) was administered to the control group. Retinal flat-mounts of each group were prepared after administration of minocycline or PBS (at age 6 weeks) and observed by laser confocal microscopy (Fig. 2). The number of Ccr2-positive cells and Cx3cr1/Ccr2 dual-positive cells were suppressed in the 50 mg/kg and 100 mg/kg minocycline-administered group compared with that in the control (Fig. 2G and H). In retina flat-mount, strict retinal layer indication is difficult. However, marked Ccr2 expression was observed at outer plexiform layer and ONL in 3D image from control group (Fig. 2D). Due to Ccr2 expression suppression by minocycline, Ccr2-RFP was hardly detected in 3D image obtained from Mino100 (Fig. 2E). By contrast, the number of Cx3cr1-positive cells was not affected by minocycline administration (Fig. 2F). The counted Cx3cr1- and Ccr2-positive cells and Cx3cr1/Ccr2 dual-positive cells are shown in percentages (Fig. 2A-C4). The proportion of Ccr2-positive and Cx3cr1/Ccr2 dual-positive cells were lower in the minocycline-administered group, indicating Ccr2 suppression by minocycline.

**Fig 2.**
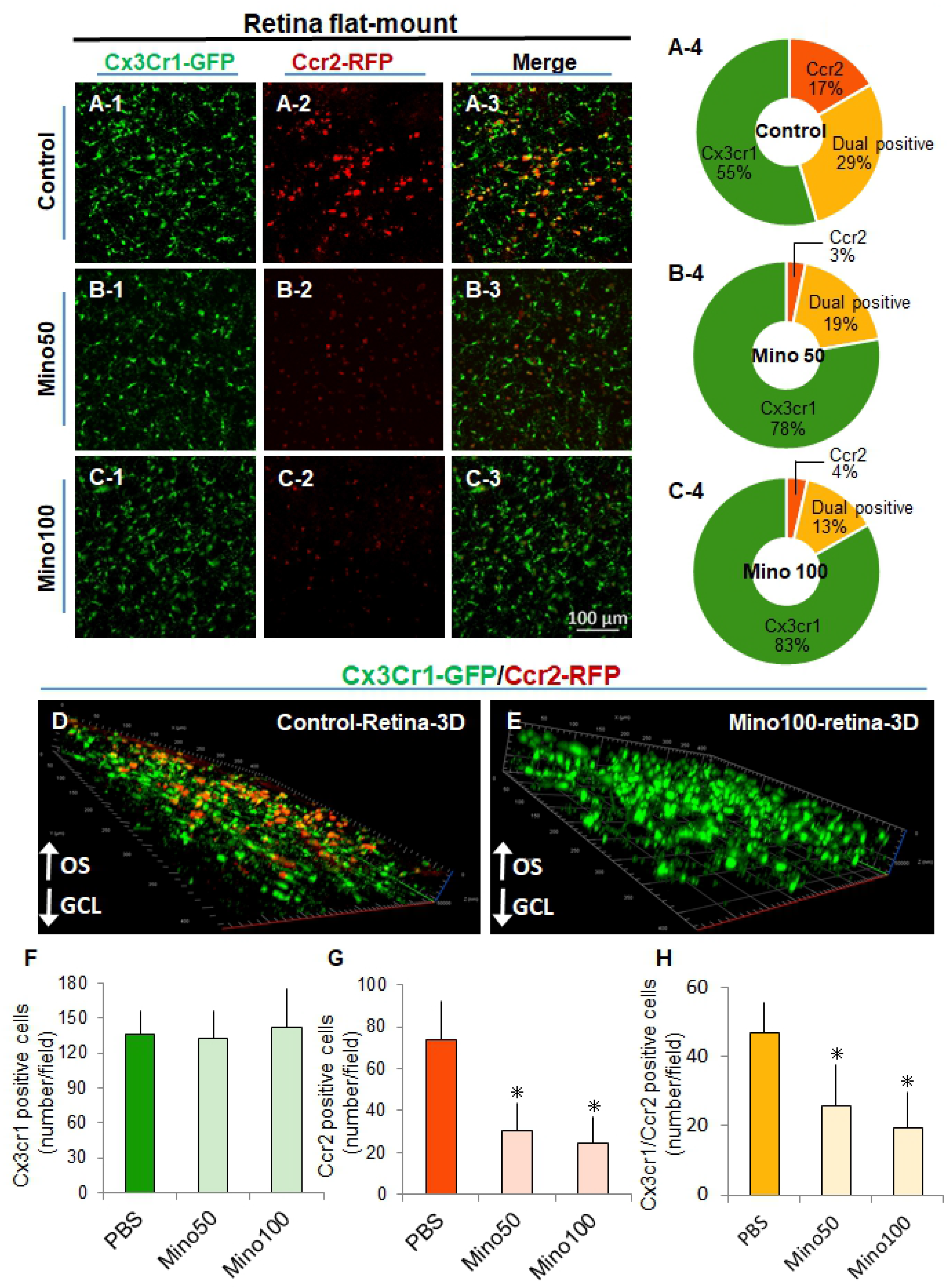
Minocycline administration suppressed Ccr2 expression in retina flat-mount. Minocycline was administered to *Mertk^-/-^Cx3cr1^GFP/+^Ccr2^RFP/+^* mice from 4-week-old to 6-week-old (continuous 14-day daily intraperitoneal [IP] injection). Minocycline group was divided into 50 mg/kg (Mino50) or 100 mg/kg (Mino100) administration. Phosphate Buffered Saline (PBS) was administered in control group. Retinal flat-mount of each group was prepared after administration of minocycline (B; Mino50, C; Mino100) or PBS (A) (at age 6 weeks) and observed by laser confocal microscopy. The percentage of Cx3cr1- and Ccr2-positive cells and Cx3cr1/Ccr2 dual-positive cells of Control, minocycline 50 mg/kg (Mino50) and 100 mg/kg (Mino100) are shown (A-4, B-4 and C-4). The 3D images from control and Mino100 are shown (D and E). The number of Cx3cr1-positive cells (D), Ccr2-positive cells (E), and Cx3cr1/Ccr2 dual-positive cells (F) from each group are shown (n ≥ 5 per group). * indicates P <.05. OS, outer segment.

### Minocycline administration suppressed Ccr2 expression in the subretinal space

RPE flat-mount was prepared to observe the apical side of RPE, corresponding to subretinal space (Fig.3) (10, 15, 18). In the 4-week-old *Mertk^-/-^Cx3cr1^GFP/+^Ccr2^RFP/+^* mice or *Mertk^+/+^Cx3cr1^GFP/+^Ccr2^RFP/+^* mice which did not show RD, neither Cx3cr1-positive cells nor Ccr2-positive cells were observed in the RPE flat-mount (data not shown) (15). Abundant Cx3cr1 expression was observed in the subretinal space in control, Mino50 and Mino100 (Fig. 3A1, B1 and C1). The number of Ccr2-positive cells and Cx3cr1/Ccr2 dual-positive cells were decreased in the 50 mg/kg (Fig.3B) and 100 mg/kg (Fig.3C) minocycline-administered group compared to the control group (Fig. 3 G, and H), though the number of Cx3cr1-positive cells did not change among groups (Fig. 3F), indicating that minocycline administration probably did not restrict migration of Cx3cr1-positive cells to subretinal space but merely suppressed Ccr2 expression. The percentages of Cx3cr1- and Ccr2-positive cells and Cx3cr1/Ccr2 dual-positive cells in RPE flat-mount are shown (Fig. 3A-C4). The proportion of Ccr2-positive and Cx3cr1/Ccr2 dual-positive cells in the subretinal space was reduced in Mino 50 and Mino 100 compared to control group.

**Fig 3.**
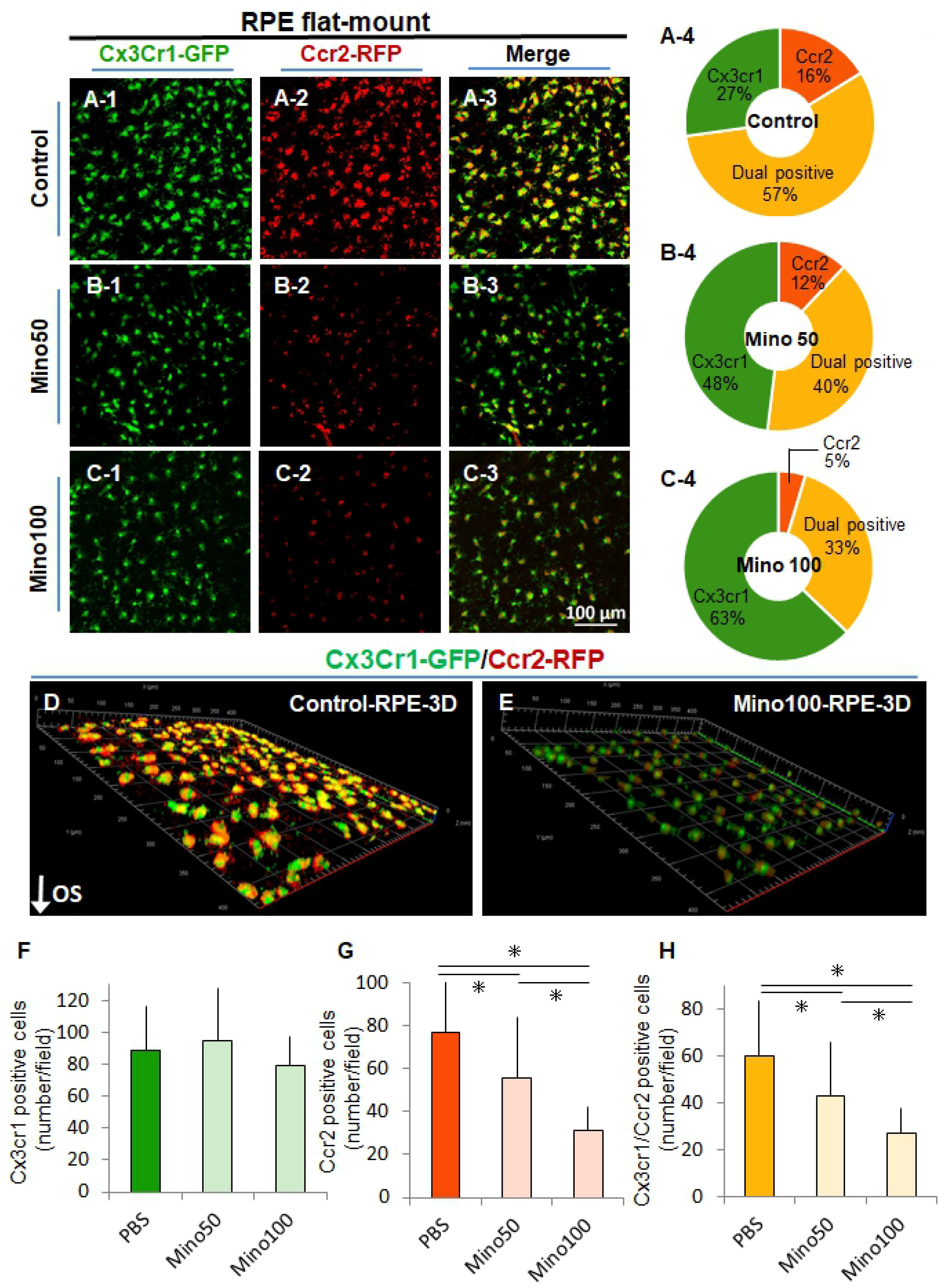
Minocycline administration suppressed Ccr2 expression in RPE flat-mount. Minocycline was administered to *Mertk^-/-^Cx3cr1^GFP/+^Ccr2^RFP/+^* mice from 4 weeks to 6 weeks of age (continuous 14-day daily intraperitoneal [IP] injection). RPE flat-mount from control (A), Mino50 (B) and Mino100 (C) was prepared after administration (at age 6 weeks). The percentage of Cx3cr1- and Ccr2-positive cells and Cx3cr1/Ccr2 dual-positive cells of Control, minocycline 50 mg/kg (Mino50) and 100 mg/kg (Mino100) are shown (A-4, B-4, C-4). The 3D images from control and Mino100 are shown (D and E). The number of Cx3cr1-positive cells (F), Ccr2-positive cells (G), and Cx3cr1/Ccr2 dual-positive cells (H) are shown (n ≥ 5 per group). * indicates P <.05.

### Amelioration of photoreceptor cell death by minocycline administration

Finally, we tested the therapeutic effect of minocycline in *Mertk^-/-^Cx3cr1^GFP/+^Ccr2^RFP/+^* mice. We have previously reported PCD amelioration by minocycline administration in a light-induced acute RD mouse model using *Abca4^-/-^Rdh8^-/-^* mice (10). However, the treatment effect of minocycline in inherited RD due to *Mertk* gene deficiency is unknown. First, minocycline was administered to 4-week old mice *Mertk^-/-^Cx3cr1^GFP/+^Ccr2^RFP/+^* mice for 2 weeks as described in Fig. 2 and 3. Nevertheless, the severity of PCD did not change between the minocycline-treated and control mice, as the PCD is relatively mild at 6 weeks of age in *Mertk^-/-^Cx3cr1^GFP/+^Ccr2^RFP/+^* mice (data not shown). Next, minocycline (50 mg/kg) or PBS was administered for 2 weeks from age 6 weeks (continuous 14-day administration) in *Mertk^-/-^Cx3cr1^GFP/+^Ccr2^RFP/+^* mice (Fig. 4). Minocycline-administered *Mertk^-/-^Cx3cr1^GFP/+^Ccr2^RFP/+^* mice showed retained outer nuclear layer (Fig. 4C) and less migrated Cx3cr1 or Ccr2-positive cells (Fig. 4A and B) compared to control group, indicating PCD amelioration by minocycline administration in inherited RD due to *Mertk* deficiency. The outer nuclear layer thickness of 4 week old *Mertk^-/-^Cx3cr1^GFP/+^Ccr2^RFP/+^* and WT (B6) mice are shown as negative control (S1 Fig).

**Figure 4.**
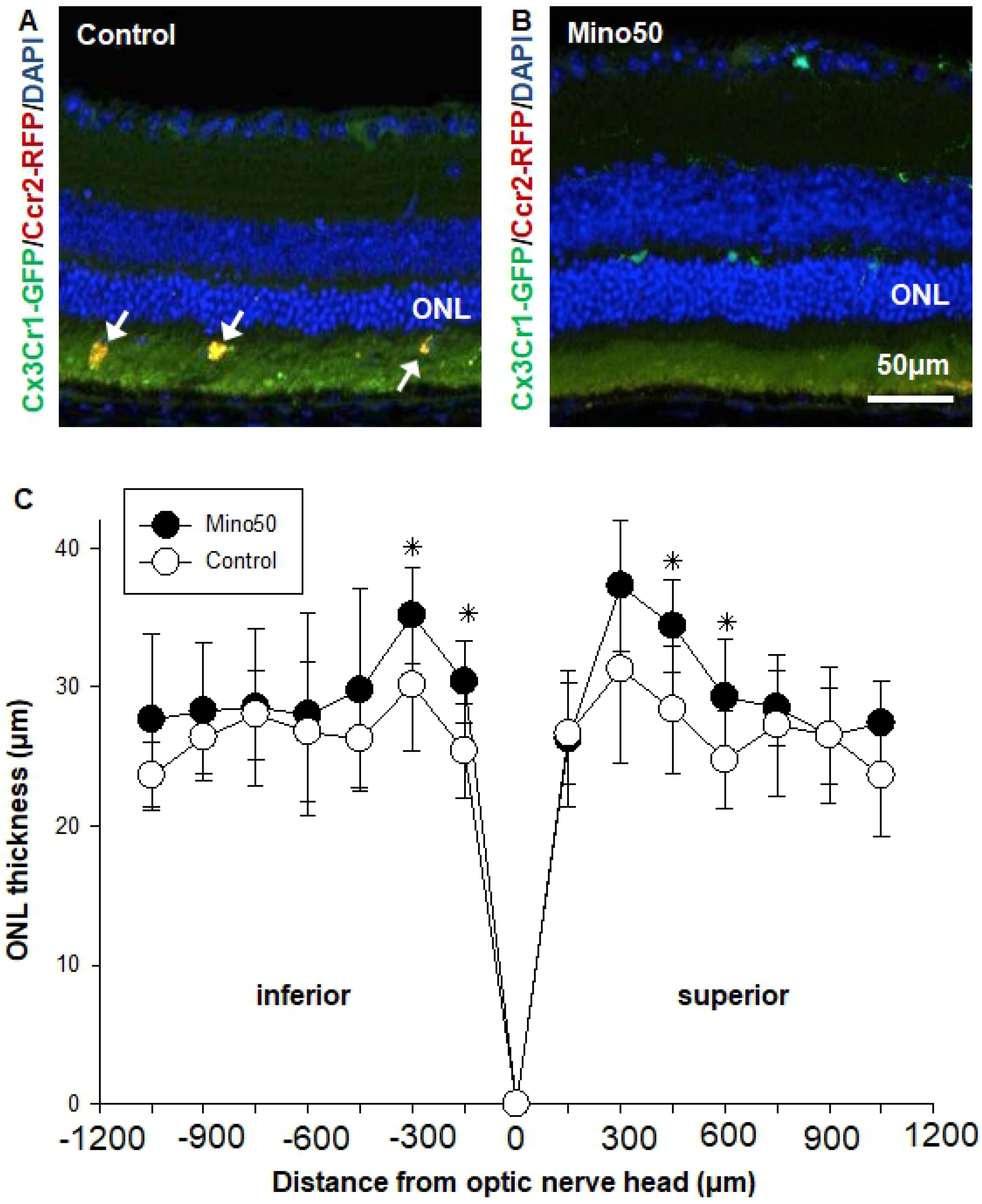
Minocycline administration ameliorates photoreceptor cell death (PCD) in *Mertk^-/-^Cx3cr1^GFP/+^Ccr2^RFP/+^* mice. Minocycline (50 mg/kg) or PBS (Control) was administered from 6 weeks to 8 weeks of age (continuous 14-day administration) *Mertk^-/-^Cx3cr1^GFP/+^Ccr2^RFP/+^* mice (C). Retinal sections of Control (A) and Mino50 (B) are shown. The thickness of outer nuclear layer of each group was measured by Image J software (C) (n ≥ 5 per group). * indicates P <.05.

## Discussion

Accumulated evidence show anti-inflammatory effect of minocycline *in vitro* and *in vivo;* including the reduction of cytokine, prostaglandin, and nitric oxide release; as well as reduced proliferation, and staining for markers such as CD11b, MHC-II and Iba-1 (7, 8, 10, 19, 20). Currently, many clinical trials are being performed for diabetic macular edema, Huntington’s disease, and multiple sclerosis (21–26). However, since minocycline is not a selective microglial inhibitor and is a semisynthetic, broad-spectrum, most lipid-soluble tetracycline of the tetracycline family; concerns of CNS side effects such as dizziness, vertigo, ataxia, and tinnitus are arising (7). Furthermore, what kind of inflammation is exactly harmful for photoreceptors is still controversial. Thus, elucidating the mechanism of action of minocycline on inflammation and photoreceptor rescue is required to discover candidate drugs for RD that are more effective with less side effects.

Recently, we developed the inherited RD model, *Mertk^-/-^Cx3cr1^GFP/+^Ccr2^RFP/+^* mouse, for visualizing expression pattern of Cx3cr1 and Ccr2 (15). Mutations in the *Mertk* gene which belongs to a family of receptor tyrosine kinases that includes AXL and TYRO3, cause retinal dystrophies in humans and animal models such as Royal College of Surgeons (RCS) rats. Cx3cr1 is the sole receptor for Cx3cl1, also called fractalkine. Cx3cr1 is expressed by dendritic cells, natural killer cells, and macrophages (27). Ccr2 is the sole receptor for Ccl2. Ccr2 is required for macrophage infiltration to injured sites (28). In CNS, Cx3cr1 but not Ccr2 is expressed in microglia from embryonic development throughout adulthood (28). In RD, Cx3cr1-positive microglia start migrating from inner retina to subretinal space (10, 15, 16, 29). In healthy retina, no Ccr2 expression is observed. We and another group have reported Ccr2 expression markedly increasing with disease progress (13, 15, 30, 31). Ccr2 positive cells are widely recognized as monocytes (30). Circulating monocytes invade the retina in degeneration ongoing stage via the retinal vessel rather than choroidal vessel, indicating breakdown of inner blood retinal barrier (13, 31). However, what is Ccr2-positive cells in subretinal space is uncertain. Abundant Ccr2- or Cx3cr1/Ccr2 dual-positive cells were observed in *Mertk^-/-^Cx3cr1^GFP/+^Ccr2^RFP/+^* mice. A recent study reported that the inflammatory cells in subretinal space are microglia-dominant and monocyte-derived macrophages are located only in outer retina (32). If this theory is applied to our model, innate immune cells in subretinal space could be divided to Cx3cr1-, Ccr2- and Cx3cr1/Ccr2 dual-positive microglia. However, the mechanism that disallowed macrophage invasion to subretinal space is not clear. The innate immune response by microglia as well as the macrophage response in RD is still a puzzle.

In the current study, similar to systemic minocycline administration in a human patient, minocycline was systemically administered to *Mertk^-/-^Cx3cr1^GFP/+^Ccr2^RFP/+^* mice. Expression pattern of Cx3cr1 and Ccr2 were observed by retinal or RPE flat-mounts, corresponding to the neural retina or subretinal space, respectively. In retinal flat-mount, the number of Cx3cr1-GFP-positive cells did not change with minocycline administration, indicating that minocycline administration does not deplete healthy microglia. This is an important and huge difference compared to microglia/macrophage depleting drugs such as clodronate liposomes or PLX5622, a colony-stimulating factor-1 receptor inhibitor (33). For elucidating the basic mechanism of microglia/macrophage participation in RD, the depleting drugs might be more powerful compared to minocycline. However, in the treatment of RD in the human patient, these depleting drugs may cause concerns regarding severe side effects. In fact, adverse effects such as accelerated weight gain, hyperactivity, and anxiolytic-like behavior were reported in PLX5622-administered juvenile mice (34). Hence, in RD, especially in RP, requiring life-long therapy, it is crucial to employ effective drugs with less side effects. From this view, minocycline has a long history of being administered as a tetracycline family member, and thus, exploring its mechanism of PCD rescue may pave way for the treatment of patients with RD. We previously insisted Ccr2 as a therapeutic target candidate for RD (15) because Ccl2 is a cognate ligand for Ccr2, whose deletion can rescue PCD in *Mertk*^-/-^ mice (18). Another group reported that deletion of Ccr2 can ameliorate RD in the model (35). However, the other group reported that although Ccl2-Ccr2 axis blockade in light exposed *Arr*^-/-^ mice reduced monocyte infiltration to retina, it did not alter the extent of retinal degeneration (13). It was discussed that this discrepancy may due to the fact that slowly progressive RD is cumulative and more affected by immune response compare to light-induced RD which PCD initiated all at once (13).

In the current study, minocycline administration not only rescued PCD but also suppressed Ccr2 expression in *Mertk^-/-^Cx3cr1^GFP/+^Ccr2^RFP/+^* mice. According to this result, Ccr2 seems like a possible therapeutic target of RD. However, it is uncertain whether direct blockade of Ccr2 can overcome the therapeutic effect of minocycline administration and thus needs to be addressed in future studies. Additionally, it should be mentioned that the role of a chemokine or its receptor might differ depending on the disease stage or be affected by the tissue environment. For example, we previously tested the role of Ccl3 (macrophage inflammatory protein 1α, MIP-1α) in RD. Ccl3 is genetically deleted in several RD models, including light-exposed and aged *Abca4^-/-^Rdh8^-/-^* and *Mertk^-/-^* mice (18). In inherited RD model, including aged *Abca4^-/-^Rdh8^-/-^* and *Mertk^-/-^* mice, Ccl3 deletion ameliorates PCD. However, in acute light-exposed *Abca4^-/-^Rdh8^-/-^* mice, PCD is exacerbated by Ccl3 deletion with unexpected increase of Ccl4 (macrophage inflammatory protein 1β, MIP-1β), which has a sequence homology of ~60% with the murine Ccl3 gene (18). The role of Ccr2 in RD should be evaluated carefully in future studies.

In summary, minocycline administration suppresses Ccr2 expression in RD model. PCD is also ameliorated by minocycline. Ccr2 suppression is one of the mechanisms of PCD rescue by minocycline.

## Supporting information

**S1 Fig. The outer nuclear layer thickness of 4-week-old *Mertk^-/-^Cx3cr1^GFP/+^Ccr2^RFP/+^* mice and WT mice.** The outer nuclear layer thickness of 4-week-old *Mertk^-/-^Cx3cr1^GFP/+^Ccr2^RFP/+^* mice and WT (B6) mice are shown as negative control of Fig.4.

